# Programmed DNA elimination of germline development genes in songbirds

**DOI:** 10.1101/444364

**Authors:** Cormac M. Kinsella, Francisco J. Ruiz-Ruano, Anne-Marie Dion-Côté, Alexander J. Charles, Toni I. Gossmann, Josefa Cabrero, Dennis Kappei, Nicola Hemmings, Mirre J. P. Simons, Juan P. M. Camacho, Wolfgang Forstmeier, Alexander Suh

## Abstract

Genomes can vary within individual organisms. Programmed DNA elimination leads to dramatic changes in genome organisation during the germline–soma differentiation of ciliates^1^, lampreys^2^, nematodes^3,4^, and various other eukaryotes^5^. A particularly remarkable example of tissue-specific genome differentiation is the germline-restricted chromosome (GRC) in the zebra finch which is consistently absent from somatic cells^6^. Although the zebra finch is an important animal model system^7^, molecular evidence from its large GRC (>150 megabases) is limited to a short intergenic region^8^ and a single mRNA^9^. Here, we combined cytogenetic, genomic, transcriptomic, and proteomic evidence to resolve the evolutionary origin and functional significance of the GRC. First, by generating tissue-specific *de-novo* linked-read genome assemblies and re-sequencing two additional germline and soma samples, we found that the GRC contains at least 115 genes which are paralogous to single-copy genes on 18 autosomes and the Z chromosome. We detected an amplification of ≥38 GRC-linked genes into high copy numbers (up to 308 copies) but, surprisingly, no enrichment of transposable elements on the GRC. Second, transcriptome and proteome data provided evidence for functional expression of GRC genes at the RNA and protein levels in testes and ovaries. Interestingly, the GRC is enriched for genes with highly expressed orthologs in chicken gonads and gene ontologies involved in female gonad development. Third, we detected evolutionary strata of GRC-linked genes. Developmental genes such as *bicc1* and *trim71* have resided on the GRC for tens of millions of years, whereas dozens have become GRC-linked very recently. The GRC is thus likely widespread in songbirds (half of all bird species) and its rapid evolution may have contributed to their diversification. Together, our results demonstrate a highly dynamic evolutionary history of the songbird GRC leading to dramatic germline–soma genome differences as a novel mechanism to minimise genetic conflict between germline and soma.

## Text

Not all cells of an organism must contain the same genome. Some eukaryotes exhibit dramatic differences between their germline and somatic genomes, resulting from programmed DNA elimination of chromosomes or fragments thereof during germline–soma differentiation^5^. Here we present the first comprehensive analyses of a germline-restricted chromosome (GRC). The zebra finch (*Taeniopygia guttata*) GRC is the largest chromosome of this songbird^6^ and likely comprises >10% of the genome (>150 megabases)^7,10^. Cytogenetic evidence suggests the GRC is inherited through the female germline, expelled late during spermatogenesis, and eliminated from the soma during early embryo development^6,11^. Previous analyses of a 19-kb intergenic region suggested that the GRC contains sequences with high similarity to regular chromosomes (‘A chromosomes’)^8^.

In order to reliably identify sequences as GRC-linked, we used a single-molecule sequencing technology not applied previously in birds that permits reconstruction of long haplotypes through linked reads^12^. We generated separate haplotype-resolved *de-novo* genome assemblies for the germline and soma of a male zebra finch (testis and liver; ‘Seewiesen’; Supplementary Table 1). We further used the linked-read data to compare read coverage and haplotype barcode data in relation to the zebra finch somatic reference genome (‘taeGut2’)^7^, allowing us to identify sequences that are shared, amplified, or unique to the germline genome in a fashion similar to recent studies on cancer aneuploidies^13^. We also re-sequenced the germline and soma from two unrelated male zebra finches (‘Spain’; testis and muscle; Extended Data Fig. 1) using short reads.

**Table 1:** The 115 high-confidence genes on the GRC with information on their A-chromosomal origin in the reference genome taeGut2, number of testis-specific SNVs, methods supporting their GRC linkage, testis/ovary RNA expression of the GRC paralog, testis/ovary protein expression of the GRC paralog, and evolutionary stratum on the GRC.

| Gene symbol | Chr. | Start | End | SNVs | Method | RNA evidence | Protein evidence | GRC stratum |
| --- | --- | --- | --- | --- | --- | --- | --- | --- |
| AAGAB | 10 | 19608548 | 19634367 | 10 | SNVs |  |  | S5 |
| ADGRL2 | 8 | 14047115 | 14171612 | 10 | SNVs |  |  |  |
| ADGRL3 | 4 | 14919933 | 15404594 | 8 | SNVs | ovary |  |  |
| AKIRIN2 | 3 | 78683482 | 78688947 | 6 | SNVs | ovary |  | S5 |
| ALDH18A1 | 6 | 36280145 | 36301392 | 17 | SNVs |  |  | S4 |
| ALG13 | 4A | 18474239 | 18501426 | 19 | SNVs | ovary |  |  |
| ARMC6 | 28 | 4942046 | 4946063 | 5 | SNVs |  |  |  |
| ATP2A2 | 15 | 2841010 | 2879975 | 8 | SNVs |  |  |  |
| BICC1 | 6 | 6355408 | 6434911 | 402 | SNVs | ovary |  | S1 |
| BMP15 | 4A | 15596686 | 15598225 | 29 | SNVs, coverage | ovary |  | S5 |
| BMPR1B | 4 | 18997710 | 19024248 | 47 | SNVs, coverage |  |  | S5 |
| CCND3 | 26_random |  |  | 14 | SNVs |  |  |  |
| CD164 | 3 | 69169111 | 69174605 | 38 | SNVs, coverage | ovary |  |  |
| COPS2 | 10 | 10200701 | 10222248 | 1 | SNVs, coverage | ovary |  |  |
| CPEB1 | 10 | 3114181 | 3137661 | 114 | SNVs | ovary |  |  |
| CSNK1A1L | Un | 135422201 | 135425792 | NA | coverage |  |  |  |
| CXCL14 | 13 | 9423543 | 9433139 | 12 | SNVs |  |  | S5 |
| DDX49 | 28 | 4913058 | 4918451 | 5 | SNVs | ovary |  |  |
| DIS3L | 10 | 19097281 | 19112154 | 13 | SNVs | ovary |  | S5 |
| DNAAF5 | 14 | 13758049 | 13780402 | NA | coverage |  |  |  |
| DNAH5 | 2 | 81235805 | 81361091 | 7 | SNVs |  |  |  |
| DPH6 | 5 | 31543945 | 31606965 | 13 | SNVs, coverage |  |  |  |
| EFNB1 | 4A | 5764021 | 5807953 | 86 | SNVs | ovary |  | S5 |
| ELAVL4 | 8 | 21034240 | 21098310 | 364 | SNVs | ovary |  |  |
| EPPK1 | Un |  |  | 52 | SNVs |  |  |  |
| FBXO16 | 3 | 112541865 | 112568948 | 6 | SNVs |  |  |  |
| FEM1B | 10 | 19886491 | 19891616 | 9 | SNVs | ovary |  | S5 |
| FIG4 | 3 | 69023384 | 69073678 | 17 | SNVs |  |  | S5 |
| FRS3 | 26_random |  |  | 42 | SNVs, coverage |  |  | S5 |
| GBE1 | 1 | 105820640 | 105934310 | 4 | SNVs, coverage |  |  |  |
| INTS9 | 3 | 112259951 | 112313512 | NA | coverage |  |  |  |
| LIAS | 4 | 48132714 | 48139736 | 42 | SNVs |  |  | S2 |
| LIN54 | 4 | 13615974 | 13637371 | 17 | SNVs |  |  |  |
| LINC02027 | 1 | 106086596 | 106087033 | NA | coverage |  |  |  |
| LMBRD2 | Z | 41646446 | 41665840 | NA | coverage |  |  |  |
| LOC100223190 | Z | 69149414 | 69156994 | 41 | SNVs |  |  |  |
| LOC100224235 | Un |  |  | 5 | SNVs |  |  | S5 |
| LOC100225322 | 1A | 47543094 | 47544622 | 6 | SNVs | ovary |  |  |
| LOC100227189 | Un | 150797142 | 150801997 | NA | coverage |  |  |  |
| LOC100228170 | Un | 55540047 | 55541360 | NA | coverage |  |  |  |
| LOC101233087 | Z | 47991391 | 47994344 | 7 | SNVs |  |  |  |
| LOC101233688 | 5 | 937818 | 939059 | 5 | SNVs |  |  | S5 |
| LOC101233767 | 18 | 8034939 | 8038005 | 11 | SNVs |  |  |  |
| LOC101233800 | Un |  |  | 16 | SNVs |  |  |  |
| LOC101234253 | 10 | 19184028 | 19186114 | 7 | SNVs | ovary |  | S5 |
| LOC105758464 | 23 | 46808 | 60360 | 14 | SNVs |  |  | S5 |
| LOC105758894 | 26_random |  |  | 5 | SNVs |  |  |  |
| LOC105758976 | 2 | 34301994 | 34306899 | 16 | SNVs |  |  |  |
| LOC105759101 | 3 | 76396180 | 76401262 | 21 | SNVs |  |  |  |
| LOC105759167 | 4A | 15573874 | 15574621 | 5 | SNVs |  |  |  |
| LOC105759195 | 4 | 14453003 | 14473747 | 18 | SNVs |  |  |  |
| LOC105759199 | 4 | 20714525 | 20720872 | 11 | SNVs |  |  |  |
| LOC105759260 | 5 | 1874731 | 1886007 | 32 | SNVs |  |  | S5 |
| LOC105759646 | Un |  |  | 7 | SNVs |  |  |  |
| LOC105759655 | Un |  |  | 8 | SNVs |  |  |  |
| LOC105759660 | Un |  |  | 18 | SNVs |  |  |  |
| LOC105759665 | Un |  |  | 5 | SNVs |  |  |  |
| LOC105759692 | Un |  |  | 12 | SNVs |  |  |  |
| LOC105759919 | Un |  |  | 8 | SNVs |  |  |  |
| LOC105760011 | Un |  |  | 7 | SNVs |  |  |  |
| LOC105760123 | Un |  |  | 18 | SNVs |  |  |  |
| LOC105760228 | Un |  |  | 14 | SNVs |  |  |  |
| LOC105760286 | Un |  |  | 18 | SNVs |  |  |  |
| LOC105760461 | Un |  |  | 10 | SNVs |  |  |  |
| LOC105760874 | Z | 60949696 | 60953194 | 19 | SNVs | testis |  |  |
| LOC105760936 | 16_random |  |  | 12 | SNVs |  |  |  |
| LUC7L3 | Un | 35019850 | 35021569 | NA | coverage |  |  |  |
| MED20 | 26_random | 110500 | 113183 | 28 | SNVs, coverage |  |  | S5 |
| MSH4 | 8 | 27964612 | 27983306 | 30 | SNVs |  |  | S4 |
| NAPA | NA |  |  | NA | Biederman et al. 2018 |  | both |  |
| NEUROG1 | 13 | 9450787 | 9451086 | 6 | SNVs |  |  |  |
| NFYA | 26 | 4725655 | 4735626 | 7 | SNVs |  |  | S5 |
| NRBP2 | 2 | 156379345 | 156398225 | 48 | SNVs |  |  |  |
| PCSK4 | 28 | 4059367 | 4063775 | 21 | SNVs |  |  |  |
| PGC | 26_random |  |  | 24 | SNVs |  |  |  |
| PHKA1 | 4A | 15562688 | 15593666 | 16 | SNVs |  |  |  |
| PIM1 | 26 | 603349 | 607242 | 50 | SNVs | testis |  |  |
| PIM3 | 1A | 18426716 | 18430551 | 81 | SNVs | ovary |  |  |
| PMM1 | 1A | 49038672 | 49047011 | NA | coverage |  |  |  |
| PRDM1 | 3 | 70624594 | 70644625 | 12 | SNVs |  |  |  |
| PRKAR1A | 18 | 2200317 | 2211579 | NA | coverage |  |  |  |
| PRKAR1B | 14 | 13784578 | 13872733 | NA | coverage |  |  |  |
| PRPSAP1 | 18 | 8008870 | 8033058 | 7 | SNVs, coverage | ovary |  | S5 |
| PSIP1 | Z | 59887174 | 59919902 | 57 | SNVs, coverage | ovary |  | S3 |
| PUF60 | 2 | 156354670 | 156376091 | 63 | SNVs | ovary |  |  |
| RFC1 | 4 | 48169638 | 48202709 | 77 | SNVs | ovary |  | S2 |
| RNF157 | 18 | 8048721 | 8062403 | NA | coverage |  |  |  |
| RNF17 | 1 | 45827734 | 45870640 | 69 | SNVs | ovary | testis | S4 |
| RNF20 | Z_random |  |  | 9 | SNVs, subtraction | both |  |  |
| ROBO1 | 1 | 107094521 | 107228509 | 19 | SNVs, coverage | ovary |  | S5 |
| ROBO2 | 1 | 107529365 | 107979302 | 25 | SNVs |  |  |  |
| RXRA | 17 | 8320685 | 8355067 | 14 | SNVs |  |  | S5 |
| SCRIB | 2 | 156239884 | 156325797 | 83 | SNVs | ovary |  | S5 |
| SECISBP2L | 10 | 10159176 | 10193647 | 60 | SNVs, coverage | ovary | both | S5 |
| SHC4 | 10 | 10124441 | 10151124 | 11 | SNVs,<br>coverage |  |  | S4 |
| SPHK1 | 18 | 7991834 | 7994408 | 2 | SNVs,<br>coverage | testis |  |  |
| SRRT | Un |  |  | 16 | SNVs | both |  |  |
| SUGP2 | 28 | 4930094 | 4937971 | 33 | SNVs | ovary | both | S5 |
| SURF4 | 17 | 7682661 | 7693000 | 50 | SNVs | ovary |  | S3 |
| TFEB | 26_rando<br>m | 20475 | 21840 | 11 | SNVs |  |  | S5 |
| TIAM2 | 3 | 54800961 | 54890499 | NA | coverage |  |  |  |
| TRIM71 | 2 | 60893878 | 60907039 | 159 | SNVs,<br>subtraction |  |  | S1 |
| UBE2O | 18 | 7960889 | 7981633 | NA | coverage |  |  |  |
| UGDH | 4 | 48113314 | 48126079 | 136 | SNVs,<br>coverage,<br>subtraction | ovary | ovary | S2 |
| UNC5C | 4 | 19035187 | 19126466 | 13 | SNVs,<br>coverage |  |  |  |
| Unnamed | Un | 12457451<br>3 | 12457555<br>3 | NA | coverage |  |  |  |
| Unnamed | Un | 12712981<br>9 | 12713050<br>3 | NA | coverage |  |  |  |
| Unnamed | 16_rando<br>m | 26580 | 73126 | NA | coverage |  |  |  |
| Unnamed | Un | 13010351<br>4 | 13010426<br>4 | NA | coverage |  |  |  |
| Unnamed | Un | 50859565 | 50860210 | NA | coverage |  |  |  |
| Unnamed | Un | 11535588<br>3 | 11535815<br>4 | NA | coverage |  |  |  |
| Unnamed | Un | 12457859<br>5 | 12457932<br>6 | NA | coverage |  |  |  |
| VEGFA | 3 | 31631385 | 31652650 | 34 | SNVs,<br>coverage | both |  |  |
| WDR19 | 4 | 48204115 | 48240398 | 34 | SNVs | ovary |  | S5 |
| ZWILCH | 10 | 19199771 | 19206407 | 8 | SNVs | ovary |  | S5 |

Note: We were able to place only some genes on evolutionary strata due to our strict criteria for evaluating the maximum likelihood gene trees. The remaining genes lacked sequence information from several of the other sampled somatic genomes or had poorly resolved tree topologies.

We first established the presence of the GRC in the three germline samples. Cytogenetic analysis using fluorescence *in-situ* hybridisation (FISH) with a new GRC probe showed that the GRC is present exclusively in the germline and eliminated during spermatogenesis as hypothesised (Fig. 1a-b, Extended Data Fig. 2)^6,11^. We compared germline/soma sequencing coverage by mapping reads from all three sampled zebra finches onto the reference genome assembly (regular ‘A chromosomes’), revealing consistently germline-increased coverage for single-copy regions, reminiscent of programmed DNA elimination of short genome fragments in lampreys^2^ (Fig. 1c-d). A total of 92 regions (41 with >10 kb length) on 13 chromosomes exhibit >4-fold increased germline coverage in ‘Seewiesen’ relative to the soma (Fig. 1e, Supplementary Table 2). Such a conservative coverage cut-off provides high confidence in true GRC-amplified regions. We obtained nearly identical confirmatory results using another library preparation method for the ‘Spain’ birds (Fig. 1f). Notably, the largest block of testis-increased coverage spans nearly 1 Mb on chromosome 1 and overlaps with the previously^8^ FISH-verified intergenic region 27L4 (Fig. 1e-f).

**Figure 1:**
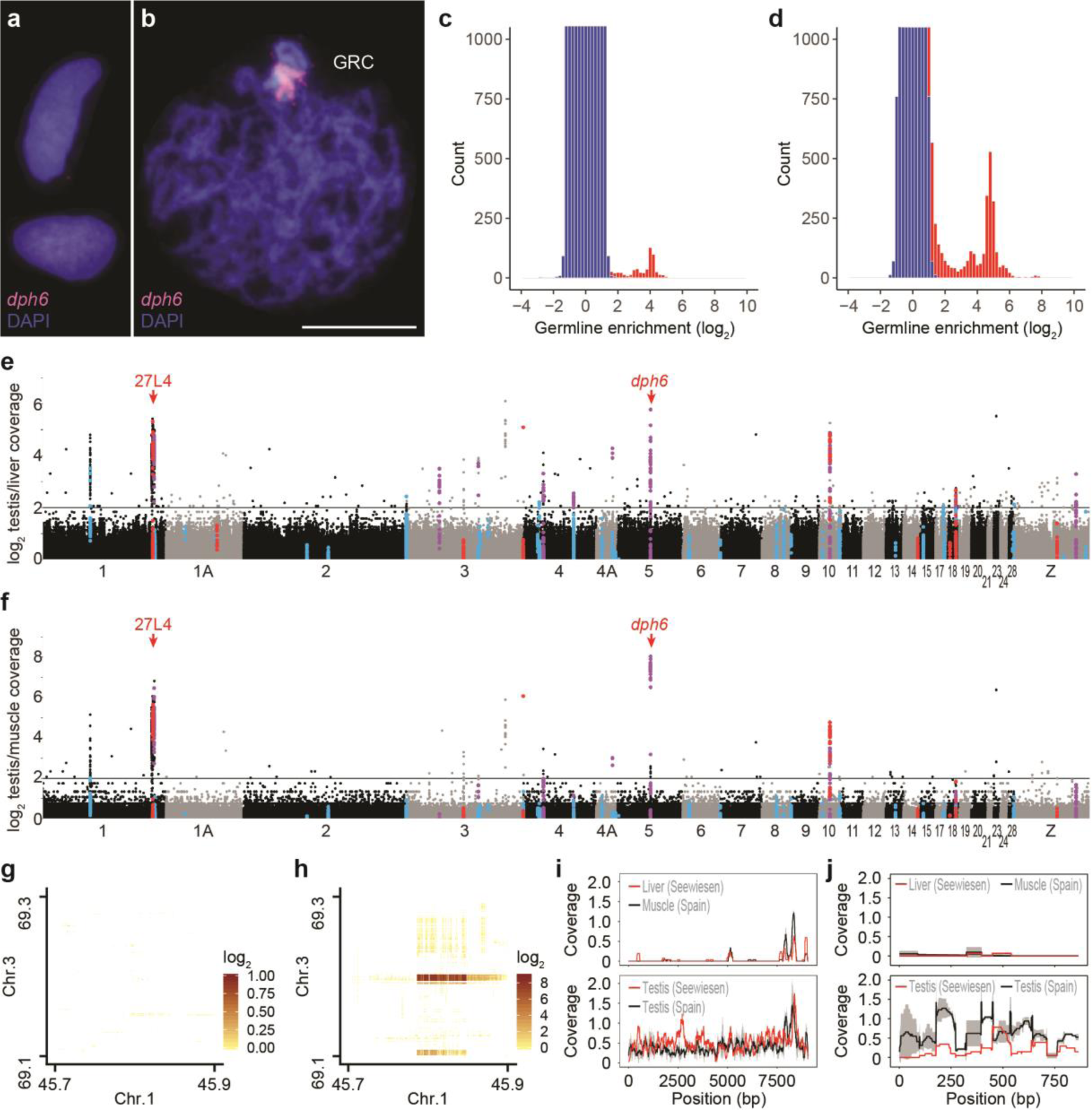
The zebra finch germline-restricted chromosome contains genes copied from many A chromosomes. **a-b**, Cytogenetic evidence for GRC absence in muscle (**a**) and GRC presence in the testis (**b**) of the same bird (Spain_1) using fluorescence *in-situ* hybridisation (FISH) of our new GRC-ampliconic probe *dph6* (selected due its high germline/soma coverage ratio; *cf*. panels e-f). The scale bar indicates 10 μm. **c-d**, Comparison of germline/soma coverage ratios (red) for 1 kb windows with an expected symmetrical distribution (blue) indicates enrichment of single-copy regions in the germline, similar to lamprey^2^ both in Spain (**c**; average of Spain_1 and Spain_2 coverage; PCR-free short reads) and Seewiesen (**d**; linked reads) samples. Y-axis is truncated for visualisation. **e-f**, Manhattan plot of germline/soma coverage ratios in 1 kb windows across chromosomes of the somatic reference genome taeGut2. Colours indicate high-confidence GRC-linked genes and their identification (red: coverage, blue: SNVs, purple: both; Table 1). Note that the similarities between Seewiesen (**e**) and Spain_1/Spain_2 averages (**f**) constitute independent biological replicates for GRC-ampliconic regions, as the data are based on different domesticated populations and different library preparation methods. Red arrows denote two FISH-verified GRC-amplified regions (*cf.* panel b)^8^. Only chromosomes >5 Mb are shown for clarity. **g-h**, Linked-read barcode interaction heatmaps of an inter-chromosomal rearrangement on the GRC absent in Seewiesen liver (**g**) but present in Seewiesen testis (**h**). **i-j**, Coverage plots of two examples of GRC-linked genes that are divergent from their A-chromosomal paralog, *trim71* (**i**) and *napa* (**j**)^9^, and thus have very low coverage (normalised by total reads and genome size) in soma.

Our linked-read and re-sequencing approach allowed us to determine the sequence content of the GRC. The GRC is effectively a non-recombining chromosome as it recombines with itself after duplication, probably to ensure stable inheritance during female meiosis^8^. We predicted that the GRC would be highly enriched in repetitive elements, similar to the female-specific avian W chromosome (repeat density >50%, compared to <10% genome-wide)^14^. Surprisingly, neither assembly-based nor read-based repeat quantifications detected a significant enrichment in transposable elements or satellite repeats in the germline samples relative to the soma samples (Extended Data Figure 3, Supplementary Table 3). Instead, most germline coverage peaks lie in single-copy regions of the reference genome overlapping 38 genes (Fig. 1e-f, Table 1, Supplementary Table 4), suggesting that these peaks stem from very similar GRC-amplified paralogs with high copy numbers (up to 308 copies per gene; Supplementary Table 5). GRC linkage of these regions is further supported by sharing of linked-read barcodes between different amplified chromosomal regions in germline but not soma (Fig. 1g-h), suggesting that these regions reside on the same haplotype (Extended Data Fig. 4). We additionally identified 245 GRC-linked genes through germline-specific single-nucleotide variants (SNVs) present in read mapping of all three germline samples onto zebra finch reference genes (up to 402 SNVs per gene; Supplementary Table 4). As a control, we used the same methodology to screen for soma-specific SNVs and found no such genes. We conservatively consider the 38 GRC-amplified genes and those with at least 5 germline-specific SNVs as our highest-confidence set (Table 1). We also identified GRC-linked genes using germline–soma assembly subtraction (Fig. 1i); however, all were already found via coverage or SNV evidence (Table 1). Together with the *napa* gene recently identified in transcriptomes (Fig. 1j)^9^, our complementary approaches yielded 115 high-confidence GRC-linked genes with paralogs located on 18 autosomes and the Z chromosome (Table 1; all 267 GRC genes in Supplementary Table 4).

We next tested whether the GRC is functional and thus probably physiologically important using transcriptomics and proteomics. We sequenced RNA from the same tissues of the two Spanish birds used for genome re-sequencing and combined these with published testis and ovary RNA-seq data from North American domesticated zebra finches^9,15^. Among the 115 high-confidence genes, 6 and 32 were transcribed in testes and ovaries, respectively (Table 1). Note, these are only genes for which we could reliably separate GRC-linked and A-chromosomal paralogs using GRC-specific SNVs in the transcripts (Fig. 2a-b, Extended Data Fig. 5, Supplementary Table 6). We next verified translation of GRC-linked genes through protein mass spectrometry data for 7 testes and 2 ovaries from another population (‘Sheffield’). From 83 genes with GRC-specific amino acid changes, we identified peptides from 5 GRC-linked genes in testes and ovaries (Fig. 2c-d, Extended Data Fig. 6, Table 1). We therefore established that many GRC-linked genes are transcribed and translated in adult male and female gonads, extending previous RNA evidence for a single gene^9^ and questioning the hypothesis from cytogenetic studies that the GRC is silenced in the male germline^16,17^. Instead, we propose that the GRC has important functions during germline development, which is supported by a significant enrichment in gene ontology terms related to reproductive developmental processes among GRC-linked genes (Fig. 2e, Supplementary Table 7). We further found that the GRC is significantly enriched in genes that are also germline-expressed in GRC-lacking species with RNA expression data available from many tissues^18^ (Fig. 2f, Supplementary Table 8). Specifically, out of 65 chicken orthologs of high-confidence GRC-linked genes, 22 and 6 are most strongly expressed in chicken testis and ovary, respectively.

**Figure 2:**
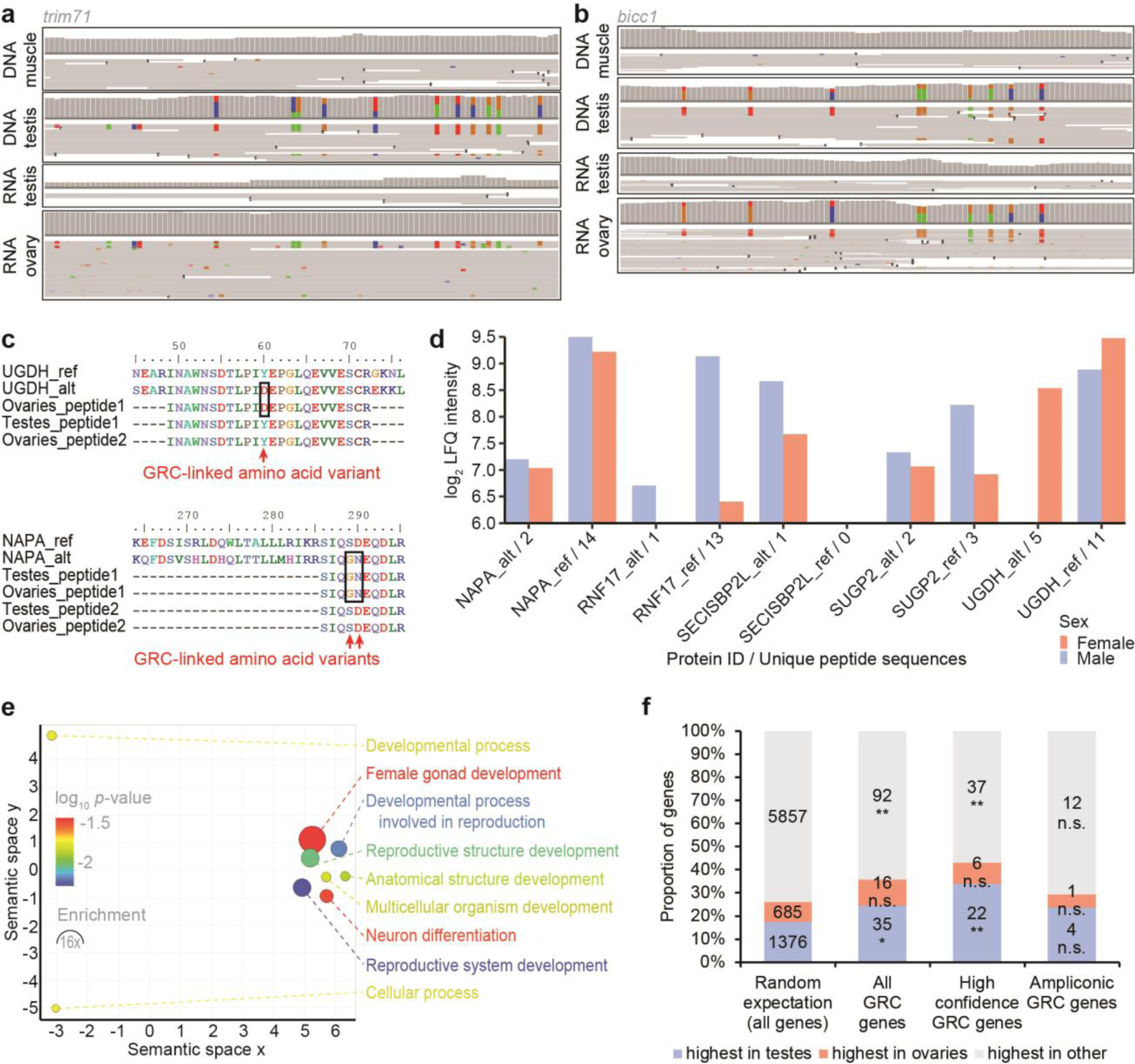
The zebra finch germline-restricted chromosome is expressed in male and female gonads. **a-b**, Comparison of coverage and read pileups for DNA-seq data from Spain_1 and Spain_2 testis/muscle, RNA-seq data from Spain_1 and Spain_2 testis, and available ovary RNA-seq data^9^. Shown are 100-bp regions within *trim71* (**a**) and *bicc1* (**b**). Colours indicate SNVs deviating from the reference genome taeGut2. **c**, Example alignments of proteomics data showing a subset of peptide expression of the respective GRC-linked paralog of *ugdh* and *napa* (alternative or ‘alt’; *cf.* reference or ‘ref’). **d**, Proteomic evidence for GRC protein expression (‘alt’) in comparison to their A-chromosomal paralog (‘ref’) of 5 genes in 7 sampled testes and 2 sampled ovaries. For label-free quantification (LFQ), unique as well as razor (non-unique) peptides were used. Note that unique peptides may occur in several of the 9 samples. **e**, Gene ontology term enrichment analysis of the 115 high-confidence GRC-linked genes (77 mapped gene symbols). Colours indicate the log_10_ of the false discovery rate-corrected *p*-value, circle sizes denote fold enrichment above expected values. **f**, Expression evidence for orthologs of three different sets of GRC genes in testes, ovaries, or other tissues in chicken^18^. Randomisation tests show a significant enrichment for germline-expressed genes among the 115 high-confidence GRC genes and all 267 GRC genes, but not the 38 ampliconic GRC genes.

The observation that all identified GRC-linked genes have A-chromosomal paralogs allowed us to decipher the evolutionary origins of the GRC. We utilised phylogenies of GRC-linked genes and their A-chromosomal paralogs to infer when these genes copied to the GRC, similarly to the inference of evolutionary strata of sex chromosome differentiation^19^. First, the phylogeny of the intergenic 27L4 locus of our germline samples and a previous GRC sequence^8^ demonstrated stable inheritance among the sampled zebra finch populations (Fig. 3a). Second, 37 gene trees of GRC-linked genes with germline-specific SNVs and available somatic genome data from other birds identify at least five evolutionary strata (Fig. 3b-f, Extended Data Fig. 7, Table 1), with all but stratum 3 containing expressed genes (*cf.* Fig. 2a-d). Stratum 1 emerged during early songbird diversification, stratum 2 before the diversification of estrildid finches, and stratum 3 within estrildid finches (Fig. 3g). The presence of at least 7 genes in these three strata implies that the GRC is tens of millions of years old and likely present across songbirds (Extended Data Fig. 7), consistent with a recent cytogenetics preprint^20^. Notably, stratum 4 is specific to the zebra finch species and stratum 5 to the Australian zebra finch subspecies (Fig. 3g), suggesting piecemeal addition of genes from 18 autosomes and the Z chromosome over millions of years of GRC evolution (Fig. 3h). The long-term residence of expressed genes on the GRC implies that they have been under selection, such as *bicc1* and *trim71* on GRC stratum 1 whose human orthologs are important for embryonic cell differentiation^21^. Using ratios of non-synonymous to synonymous substitutions (dN/dS) for GRC-linked genes with >50 GRC-specific SNVs, we found 17 genes evolving faster than their A-chromosomal paralogs (Supplementary Table 9). However, we also detected long-term purifying selection on 9 GRC-linked genes, including *bicc1* and *trim71*, as well as evidence for positive selection on *puf60*, again implying that the GRC is an important chromosome with a long evolutionary history.

**Figure 3:**
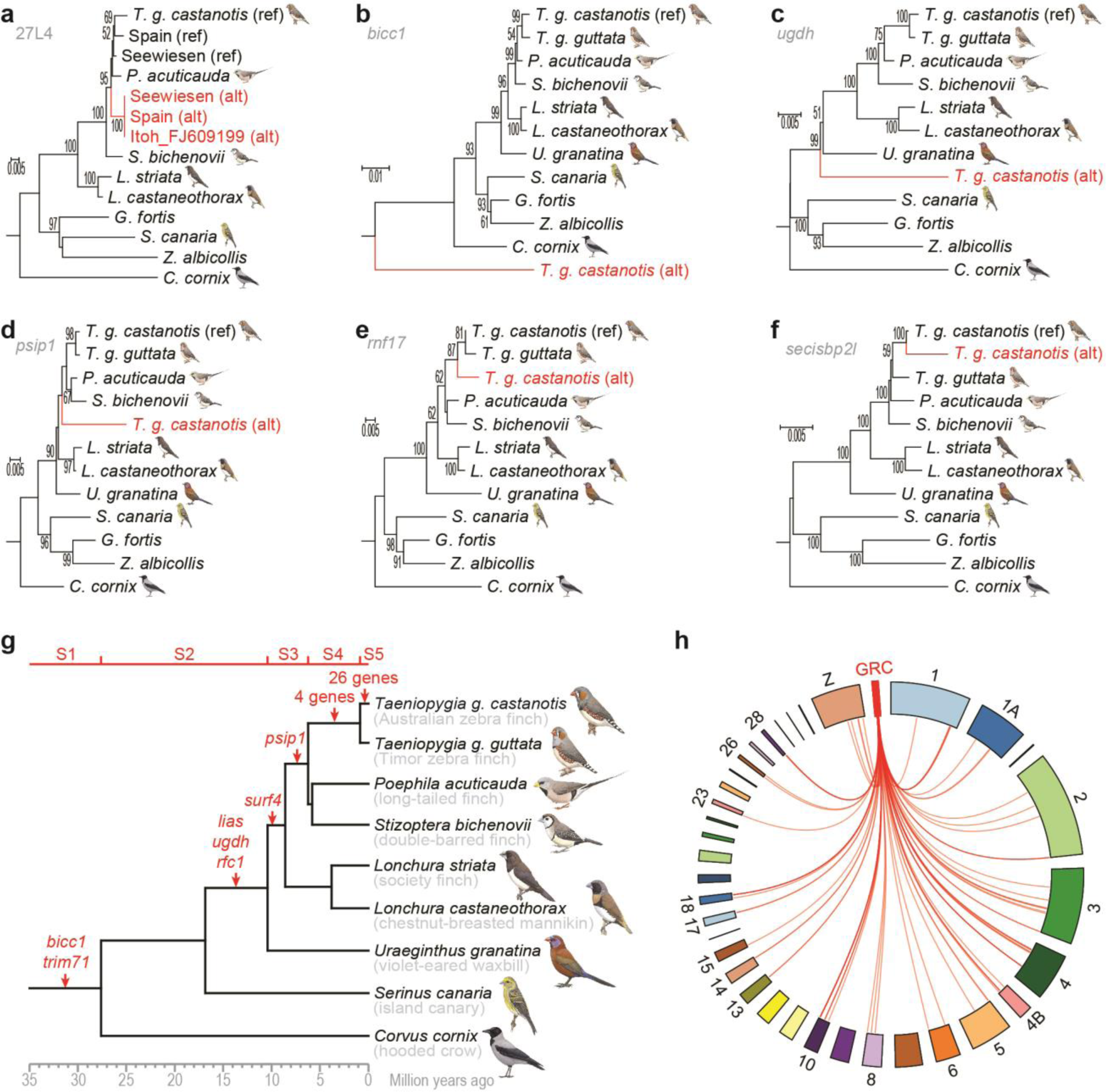
The zebra finch germline-restricted chromosome is ancient and highly dynamic. **a**, Phylogeny of the intergenic 27L4 locus previously sequenced by Itoh et al.^8^ suggests stable inheritance of the GRC paralog (alternative or ‘alt’ in red; *cf.* reference or ‘ref’) among the sampled zebra finches. **b-f**, Phylogenies of GRC-linked genes (‘alt’, in red; most selected from expressed genes) diverging from their A-chromosomal paralogs (‘ref’) before/during early songbird evolution (**b**; *bicc1*, stratum 1; *cf.* Extended Data Fig. 7), during songbird evolution (**c**; *ugdh*, stratum 2), during estrildid finch evolution (**d**; *psip1*, stratum 3), in the ancestor of the zebra finch species (**e**; *rnf17*, stratum 4), and in the Australian zebra finch subspecies (**f**; *secisbp2l*; stratum 5). The maximum likelihood phylogenies in panels a-f (only bootstrap values ≥50% shown) include available somatic genome data from estrildid finches and other songbirds. **g**, Species tree of selected songbirds showing the emergence of evolutionary strata (S1– S5) on the GRC (red gene names). Molecular dates are based on previous phylogenies^22,27^. Bird illustrations were used with permission from Lynx Edicions. **h**, Circos plot indicating A-chromosomal origin of high-confidence GRC-linked genes from 18 autosomes and the Z chromosome. Note that A-chromosomal paralogs of 37 genes remain unplaced on chromosomes in the current zebra finch reference genome taeGut2.

Here we provided the first evidence for the origin and functional significance of a GRC. Notably, our analyses suggest that the GRC emerged during early songbird evolution and we predict it to be present in half of all bird species. The species-specific addition of dozens of genes on stratum 5 implies that the rapidly evolving GRC likely contributed to reproductive isolation during the massive diversification of songbirds^22^. It was previously hypothesised that GRCs are formerly parasitic B chromosomes that became stably inherited^23,24^. Our evidence for an enrichment of germline-expressed genes on the zebra finch GRC is reminiscent of nematodes and lampreys where short genome fragments containing similar genes are eliminated during germline–soma differentiation^2–4^. All these cases constitute extreme mechanisms of gene regulation through germline–soma gene removal rather than transcriptional repression^3,5,10^. Remarkably, the GRC harbours several genes involved in the control of cell division and germline determination, including *prdm1*, a key regulator of primordial germ cell differentiation in mice^25,26^. Consequently, we hypothesise that the GRC became indispensable for its host by the acquisition of germline development genes and probably acts as a germline-determining chromosome. The aggregation of developmental genes on a single eliminated chromosome constitutes a novel mechanism to ensure germline-specific gene expression in multicellular organisms. This may allow adaptation to germline-specific functions free of detrimental effects on the soma which would otherwise arise from antagonistic pleiotropy.

## Acknowledgements

We thank Peter Ellis, Moritz Hertel, Martin Irestedt, Regine Jahn, Max Käller, Bart Kempenaers, Ulrich Knief, Pedro Lanzas, Juan Gabriel Martínez, Julio Mendo-Hernández, Beatriz Navarro-Domínguez, Remi-André Olsen, Mattias Ormestad, Yifan Pei, Douglas Scofield, Linnéa Smeds, Venkat Talla, and members of the Barbash lab and the Suh lab for support and discussions. Mozes Blom, Jesper Boman, Nazeefa Fatima, James Galbraith, Octavio Palacios, and Matthias Weissensteiner provided helpful comments on an earlier version of this manuscript. A.S. was supported by grants from the Swedish Research Council Formas (2017-01597), the Swedish Research Council Vetenskapsrådet (2016-05139), and the SciLifeLab Swedish Biodiversity Program (2015-R14). The Swedish Biodiversity Program has been made available by support from the Knut and Alice Wallenberg Foundation. F.J.R.R., J.C., and J.P.M.C. were supported by the Spanish Secretaría de Estado de Investigación, Desarrollo e Innovación (CGL2015-70750-P), including FEDER funds, and F.J.R.R. was also supported by a Junta de Andalucía fellowship. A.M.D.C was supported by a postdoc fellowship from Sven och Lilly Lawskis fond. T.I.G. was supported by a Leverhulme Early Career Fellowship Grant (ECF-2015-453). T.I.G., A.J.C. (CABM DTP), and M.S. were supported by a NERC grant (NE/N013832/1). N.H. was supported by a Patrick & Irwin-Packington Fellowship from the University of Sheffield and a Royal Society Dorothy Hodgkin Fellowship. D.K. was supported by the National Research Foundation Singapore and the Singapore Ministry of Education under its Research Centres of Excellence initiative. W.F. was supported by the Max Planck Society. Some of the computations were performed on resources provided by the Swedish National Infrastructure for Computing (SNIC) through Uppsala Multidisciplinary Center for Advanced Computational Science (UPPMAX). The authors acknowledge support from the National Genomics Infrastructure in Stockholm funded by Science for Life Laboratory, the Knut and Alice Wallenberg Foundation and the Swedish Research Council.

## Author Contributions

Conceptualisation: W.F., A.S., J.P.M.C., F.J.R.R., C.M.K., A.M.D.C., T.I.G.; cytogenetics analyses and interpretation: J.P.M.C., F.J.R.R., J.C.; genomic analyses and interpretation: A.S., C.M.K., F.J.R.R., A.M.D.C.; transcriptomic analyses and interpretation: F.J.R.R.; proteomic analyses and interpretation: T.I.G., A.J.C., D.K., M.J.P.S., N.H.; gene enrichment analyses and interpretation: C.M.K., W.F., A.S.; phylogenetic analyses and interpretation: F.J.R.R., A.S., C.M.K., T.I.G.; manuscript writing: A.S. with input from all authors; methods and supplements writing: C.M.K. with input from all authors; supervision: A.S., J.P.M.C., T.I.G., M.J.P.S. All authors read and approved the manuscript.

## Author Information

The authors declare no competing financial interests. Correspondence and requests for materials should be addressed to F.J.R.R. and A.S.

---

**Extended Data Figure 1:**
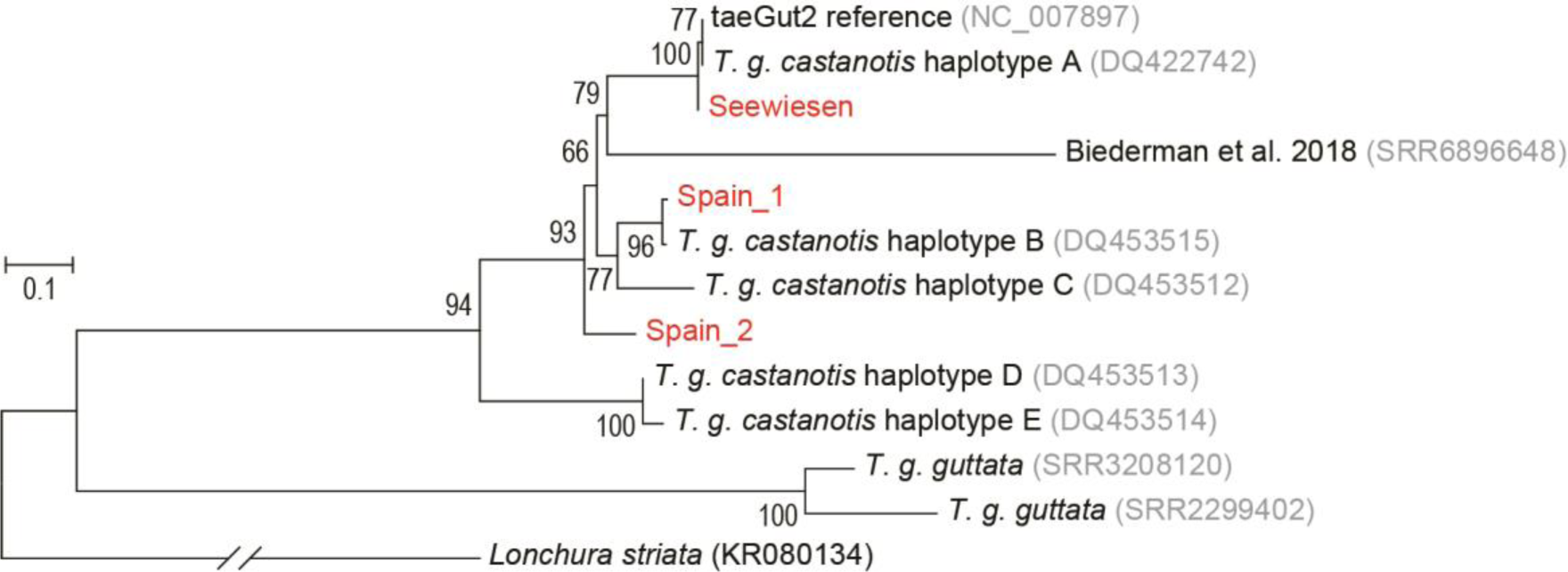
Maximum likelihood phylogeny of the five zebra finch mitochondrial haplotypes described by Mossman et al.^28^ and mitogenomes assembled from all zebra finch Illumina libraries used in this work, comprising both the Australian zebra finch (*Taeniopygia guttata castanotis*) and the Timor zebra finch (*Taeniopygia guttata guttata*) subspecies. Note that the three individuals sequenced by us (red colour) and by Biederman et al.^9^ belong to different mitochondrial haplotypes.

**Extended Data Figure 2:**
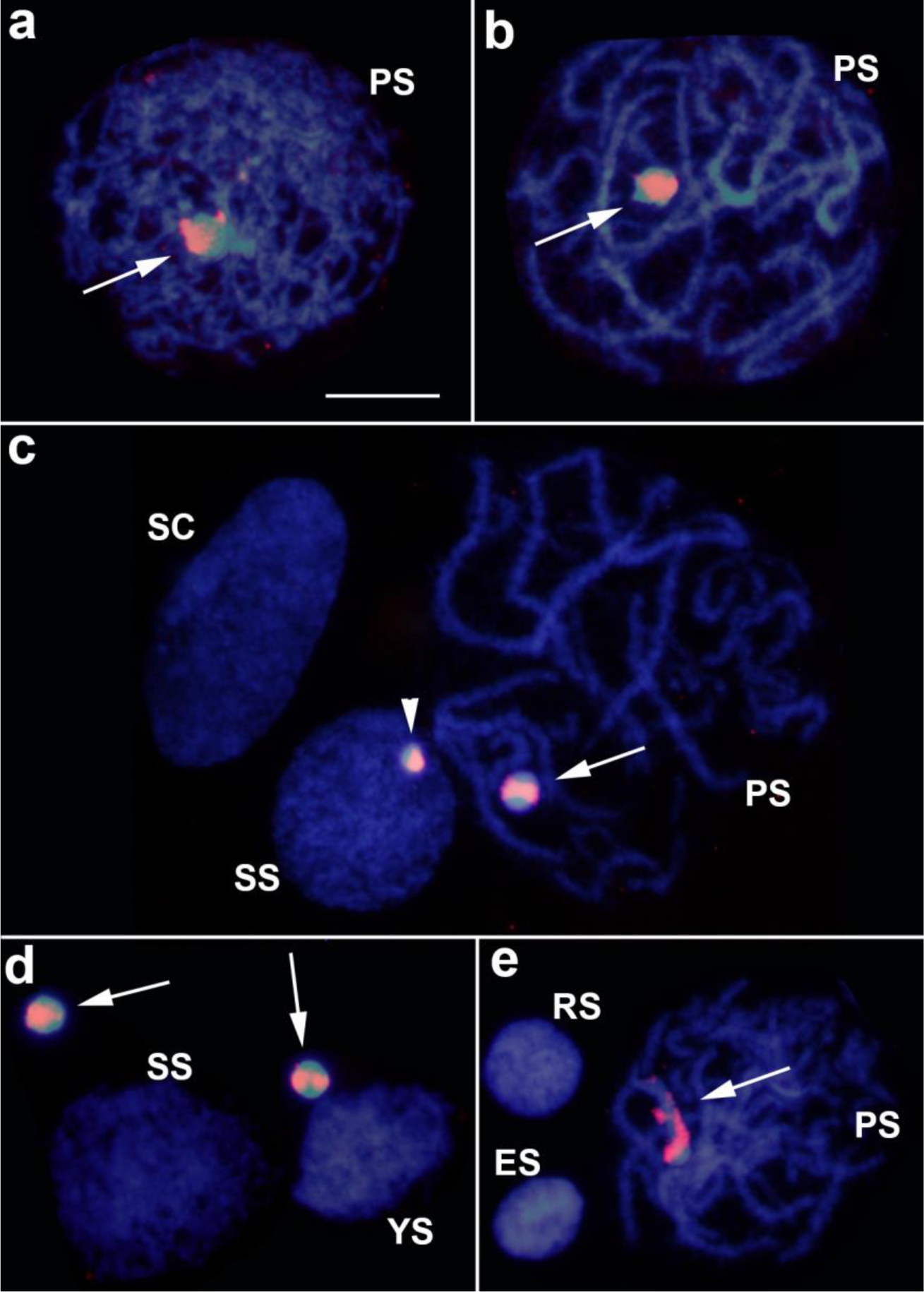
FISH analysis in testis cells of the Spain_1 zebra finch individual using the *dph6* probe (red) counterstained with DAPI (blue). Note the presence of primary (PS) and secondary (SS) spermatocytes, young spermatids (YS) and maturing spermatids at round (RS) and elongating (ES) stages. Also note that the *dph6* probe hybridises with only part of the GRC chromosome (arrow), and this is apparent in PS at leptotene-zygotene (**a**), pachytene (**b-c**, **e**) and in GRCs which failed to integrate into the main nucleus of SS or YS cells (**d**), with no FISH signal in somatic cells (SC) indicating GRC absence in somatic structural testis cells (**c**). The half size of GRC in the SS cell in panel c, compared with that in the PS next to it and that those lying outside nuclei in panel d, indicates that GRC sometimes divides equationally in the first meiotic division (resulting in the half sized GRC body in panel c) but, in most cases, it divides reductionally yielding the large sized GRCs in panel d. Note that RS and ES nuclei in panel e lack FISH signal, indicating GRC absence. All photographs were made at the same magnification, and the scale bar in panel a indicates 10 μm.

**Extended Data Figure 3:**
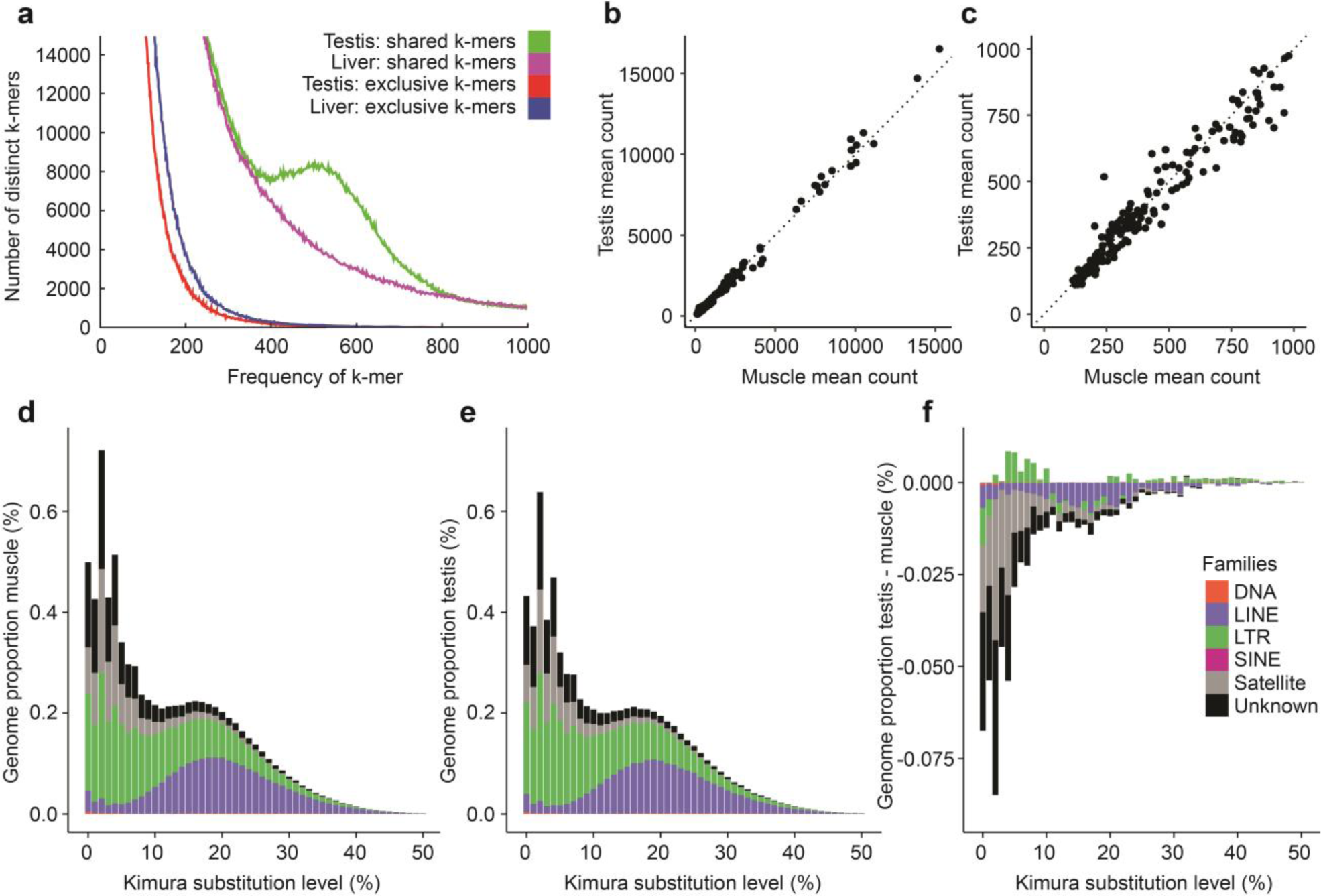
The zebra finch GRC is not enriched in satellites or specific transposable element families. **a**, Comparison of spectra for k-mers shared between or exclusive to genome sequencing data from testis and liver of the Seewiesen sample, showing that the germline is not enriched for exclusive high frequency k-mers, but is conspicuously enriched in high frequency k-mers shared with the soma. **b**, Comparison of simple repeat abundance as assessed by kSeek in the Spanish muscle samples relative to the testis samples. **c**, Same as in panel b, with a focus on low abundance simple repeats. **d-e**, Repeat landscapes based on RepeatMasker analyses showing the main repetitive element families for genome re-sequencing data from muscle (**d**) and testis (**e**) of the combined Spanish samples. **f**, Subtractive repeat landscape obtained by subtracting muscle from testis counts showing a general impoverishment of testis for most of the repetitive elements (negative values) due to the presence of the GRC.

**Extended Data Figure 4:**
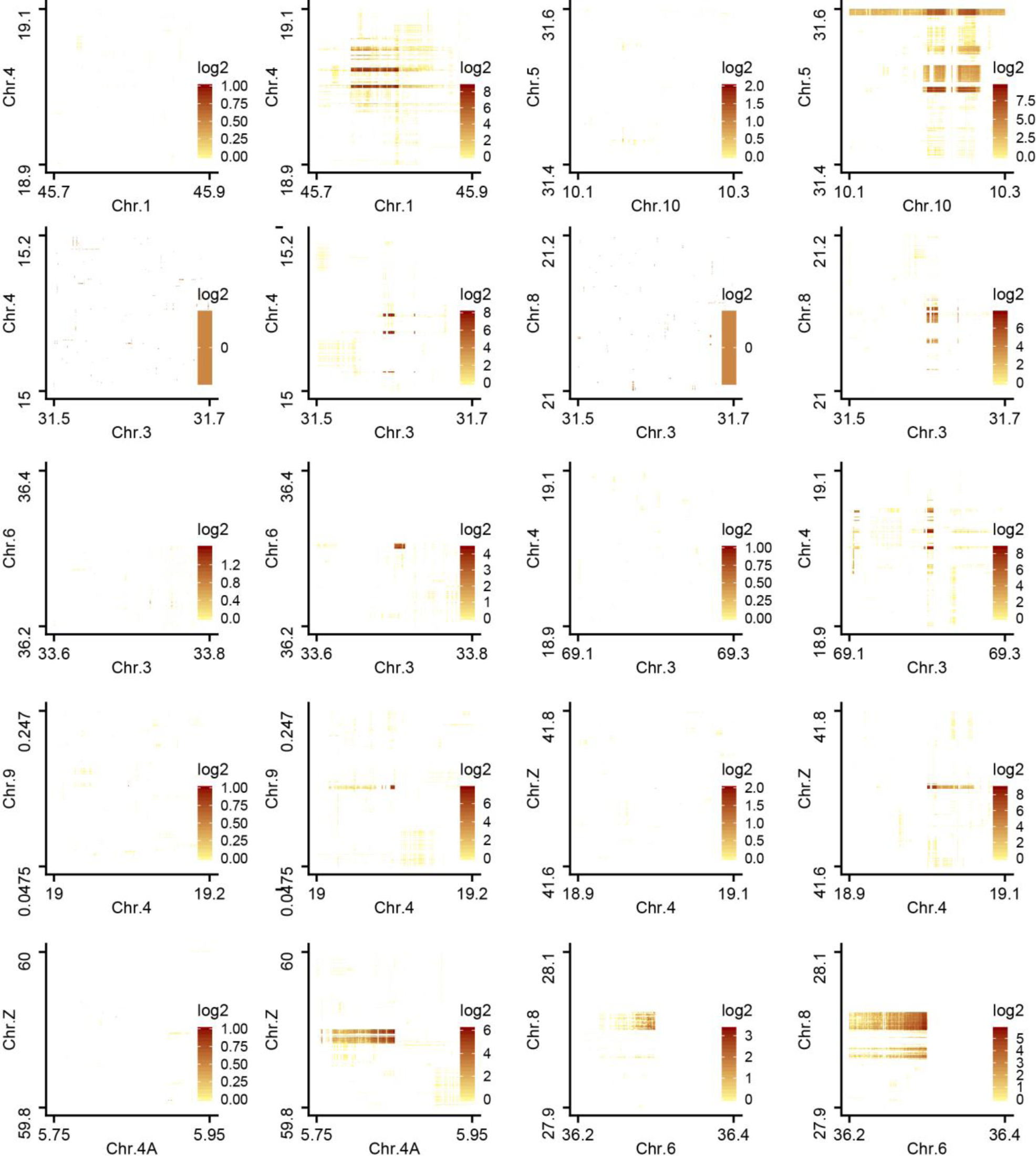
Testis-specific linked-read barcode sharing between A chromosomes indicates GRC haplotypes. Plots show side-by-side comparison of the inter-chromosomal barcode overlap for 200-kb regions for the liver and testis, respectively (chromosome position scale in Mb). With the exception of the interaction between chromosome 6 and chromosome 8 (bottom right) showing some background in the liver sample (potentially due to a shared A-chromosomal rearrangement), all inter-chromosomal structural variants were testis-specific and thus indicative of being on the same haplotype on the GRC. We exported barcode overlap matrices from the Loupe browser for testis-specific structural variants called by LongRanger and plotted them in R (v. 3.5.1). We reassigned 0 values to “NA” (shown in white on the plot) and log_2_-transformed all values. Note that the scale varies across plots.

**Extended Data Figure 5:**
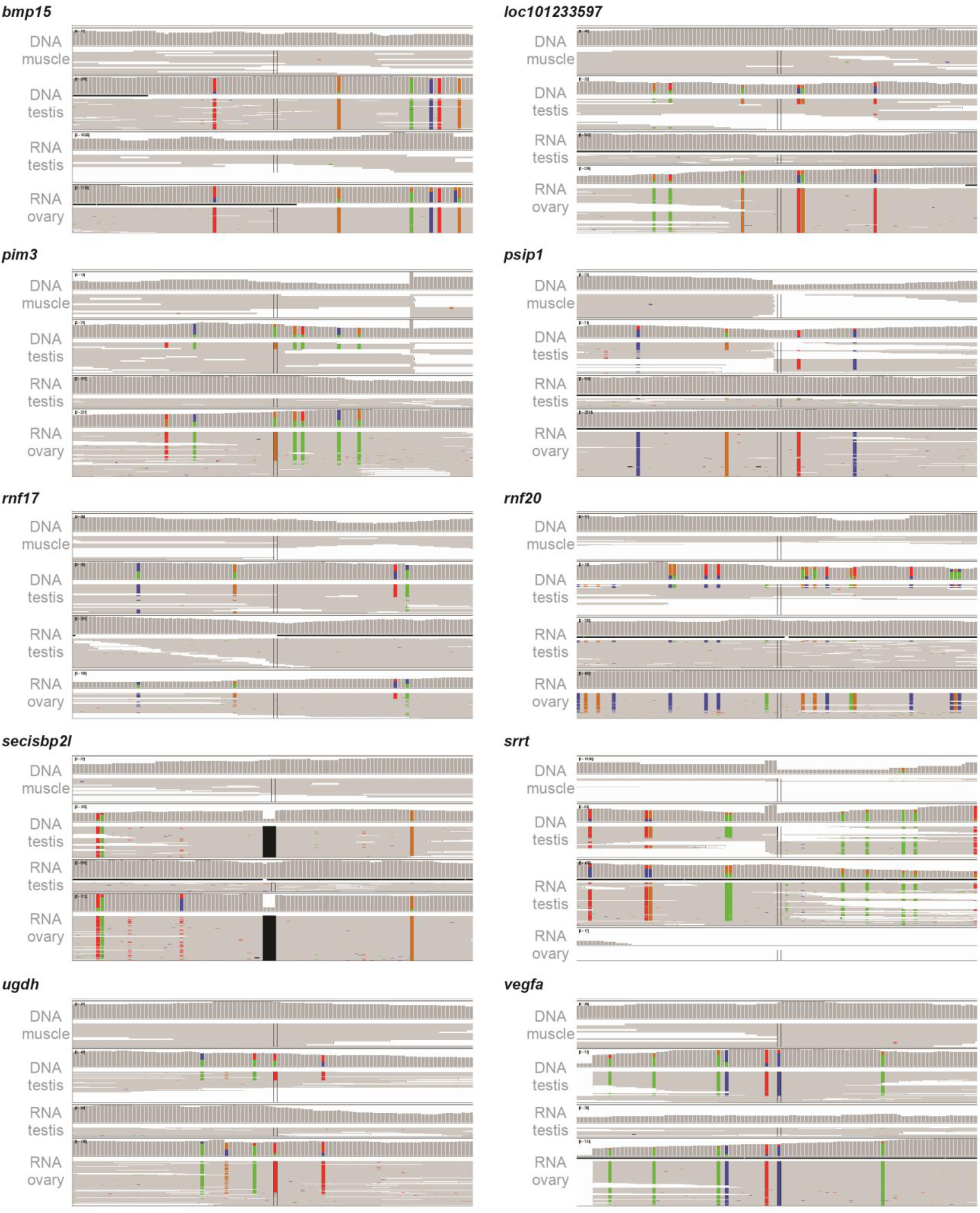
Further examples for RNA expression of GRC-linked genes. Comparison of coverage and read pileups for DNA-seq from Spain_1 and Spain_2 testis/muscle, RNA-seq data from Spain_1 and Spain_2 testis, and available ovary RNA-seq data^9^. Shown are 100-bp regions within 10 selected genes. Colours indicate SNVs deviating from the zebra finch reference genome taeGut2.

**Extended Data Figure 6:**
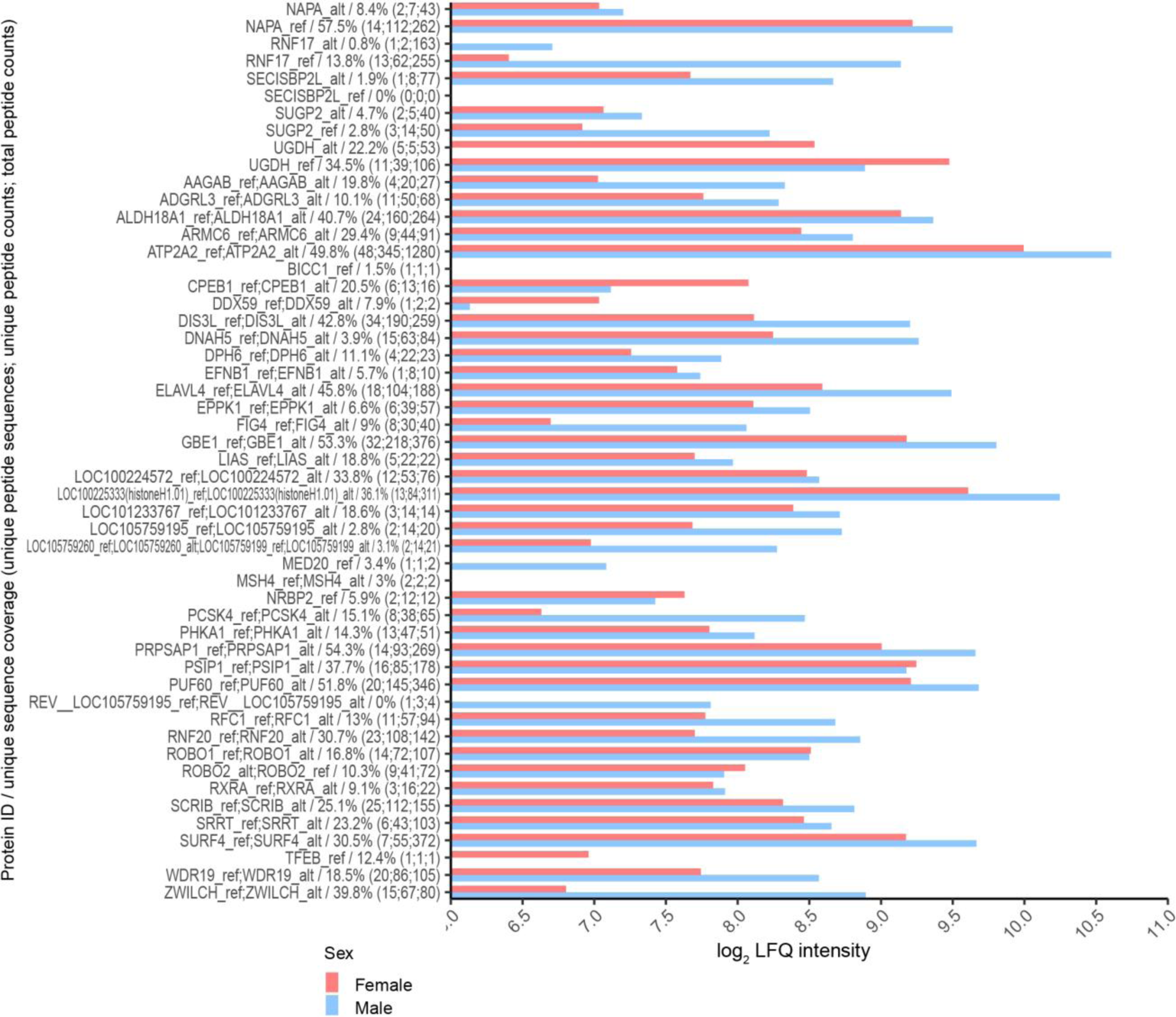
Proteomic evidence for GRC protein presence in zebra finch testes and ovaries. The five proteins listed at the top are also shown in Fig. 2d, i.e., those where we could differentiate between peptides from GRC vs. A chromosomes. GRC paralogs are denoted by the ‘alt’ suffix, whereas A-chromosomal paralogs are denoted by the ‘ref’ suffix. Unique sequence coverage corresponds to the peptide coverage percentage of the reference protein sequence. Note that unique peptides may occur in several samples (testes/ovaries). Entries of only one protein identification have sufficient evidence at the peptide level to differentiate between the GRC and A-chromosomal paralogs due to coverage of non-identical regions between the both reference sequences; entries of more than one protein identification contain evidence of presence based solely on identical regions, thus cannot be differentiated at the proteomic level. Entries of only one protein identification without the corresponding ‘alt’ or ‘ref’ variant contain evidence that span the non-identical region only, thus the alternate variant need not be called.

**Extended Data Figure 7:**
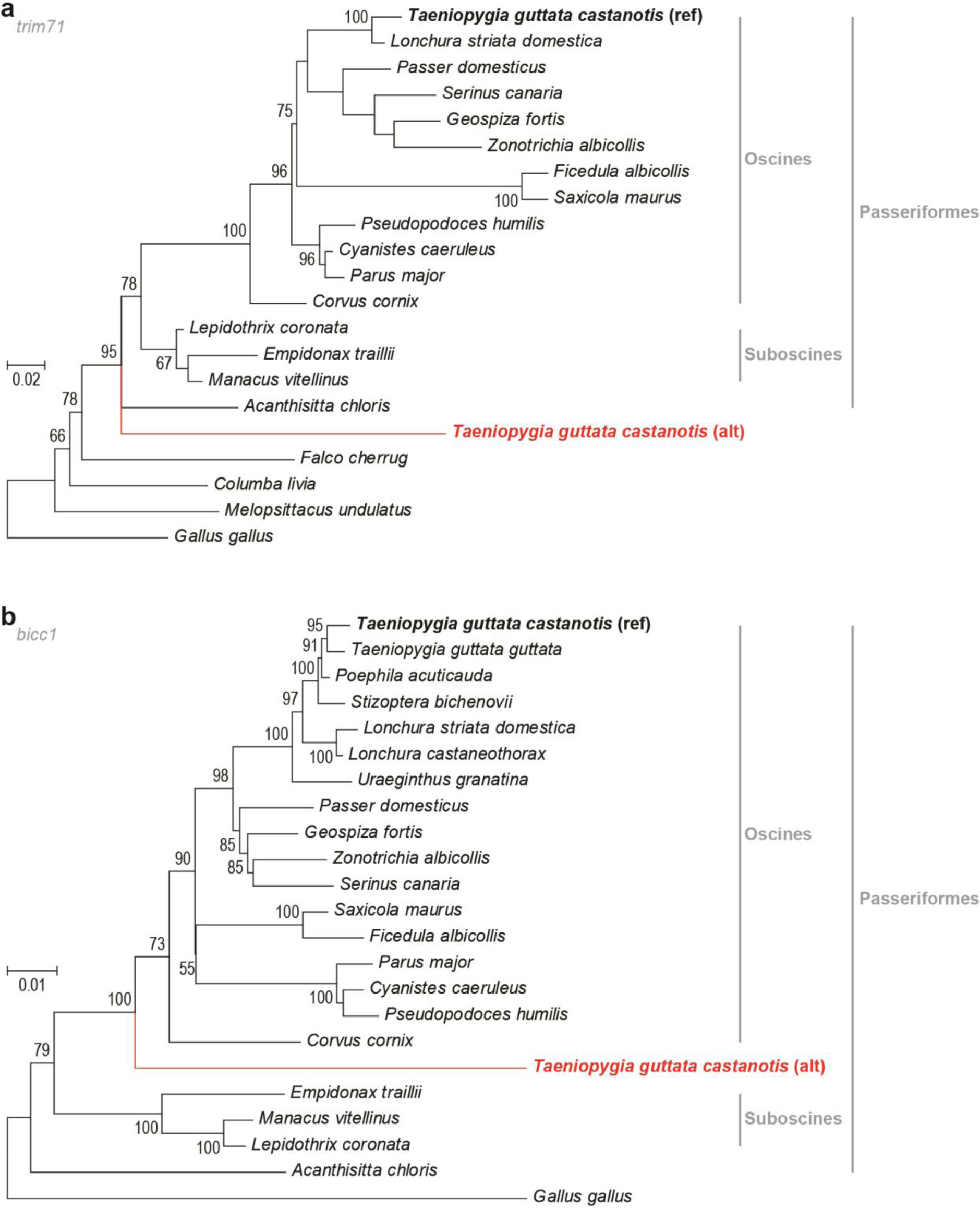
Gene trees of GRC-linked genes from stratum 1 and their A-chromosomal paralogs from broad taxon sampling imply GRC emergence in the ancestor of Passeriformes. **a**, Maximum likelihood gene tree of *trim71* (partitioned for codon positions) suggesting GRC linkage in the ancestor of Passeriformes. **b**, Maximum likelihood gene tree of *bicc1* (only 3’ UTR) suggesting GRC linkage in the ancestor of oscine songbirds.

## Supplementary Information

### Methods and Supplementary Text

**Supplementary Table 1 | Assembly metrics of linked-read *de-novo* assemblies generated from liver and testis samples of the Seewiesen zebra finch individual.**

**Supplementary Table 2 | Position, length, and library source of genomic blocks >10-kb with average germline/soma corrected coverage >4, with respect to the zebra finch reference genome (taeGut2).**

**Supplementary Table 3 | Repeat annotation of the pseudohaploid testis and liver *de-novo* assemblies from the Seewiesen zebra finch individual.**

**Supplementary Table 4 | All 267 genes on the GRC with information on their A-chromosomal origin in taeGut2, number of testis-specific SNVs, methods supporting their GRC linkage, testis/ovary RNA expression of the GRC paralog, testis/ovary protein expression of the GRC paralog, and evolutionary stratum on the GRC.**

**Supplementary Table 5 | Copy number estimates for 61 GRC-linked genes with at least 2 copies on the GRC as estimated from excess coverage in testis.**

**Supplementary Table 6 | Transcriptome analyses of GRC-linked genes showing the number of ‘alt’ SNVs per transcript with a minimum of 100 reads and an ‘alt’/’ref’ SNV mapping ratio above 1% in testes and ovary RNA-seq data.**

**Supplementary Table 7 | Enriched gene ontology terms for 167 mapped gene symbols from all 267 GRC-linked genes, and 77 mapped genes from 115 high confidence genes.**

**Supplementary Table 8 | Enrichment analyses of GRC gene orthologs in chicken and human RNA-seq data for testes, ovaries, and other tissues.**

**Supplementary Table 9 | Codon substitution rate analyses for 17 genes with at least 50 GRC-specific SNVs.**

**Supplementary Data**

